# Older age at Initiation of Antiretroviral Therapy Predicts Low Bone Mineral Density in Children with perinatally-infected HIV in Zimbabwe

**DOI:** 10.1101/565143

**Authors:** Celia L Gregson, April Hartley, Edith Majonga, Grace Mchugh, Nicola Crabtree, Ruramayi Rukuni, Tsitsi Bandason, Cynthia Mukwasi-Kahari, Kate A Ward, Hilda Mujuru, Rashida A Ferrand

## Abstract

**Background:** Perinatally-acquired HIV infection commonly causes stunting in children, but how this affects bone and muscle development is unclear. We investigated differences in bone and muscle mass and muscle function between children with HIV (CWH) and uninfected children.

**Setting:** Cross-sectional study of CWH (6–16 years) receiving antiretroviral therapy (ART) for >6 months and children in the same age-group testing HIV-negative at primary health clinics in Zimbabwe.

**Methods:** From Dual-energy X-ray Absorptiometry (DXA) we calculated total-body less-head (TBLH) Bone Mineral Content (BMC) for lean mass adjusted-for-height (TBLH-BMC^LBM^) Z-scores, and lumbar spine (LS) Bone Mineral Apparent Density (BMAD) Z-scores.

**Results:** The 97 CWH were older (mean age 12.7 *vs*. 10.0 years) and therefore taller (mean height 142cm *vs*. 134cm) than those 77 uninfected. However, stunting (height-for-age Z-score≤-2) was more prevalent in CWH (35% *vs*. 5%, *p*<0.001). Amongst CWH, 15% had low LS-BMAD (Z-score ≤-2) and 13% had low TBLH-BMC^LBM^, vs. 1% and 3% respectively in those uninfected (both *p*≤0.02). After age, sex, height and puberty adjustment, LS-BMAD was 0.33 SDs (95%CI −0.01, 0.67; *p*=0.06) lower in CWH, with no differences in TBLH-BMC^LBM^, lean mass or grip strength by HIV status. However, there was a strong relationship between age at ART initiation and both LS-BMAD Z-score (r=-0.33, *p*=0.001) and TBLH-BMC^LBM^ Z-score (r=-0.23, *p*=0.027); for each year ART initiation was delayed a 0.13 SD reduction in LS-BMAD was seen.

**Conclusion:** Size-adjusted low bone density is common in CWH. Delay in initiating ART adversely affects bone density. Findings support immediate ART initiation at HIV diagnosis.

## Introduction

In 2017, globally, up to 1.8 million children were living with HIV^1^, 90% in sub-Saharan Africa where HIV remains the leading cause of death^2^. The scale-up of antiretroviral therapy (ART) has dramatically improved survival for children living with HIV^3^; changing the infection from one that was almost invariably fatal to a chronic treatable, but incurable, condition. However, even in the era of ART, children experience a range of multisystem morbidities due to their infection and/or their treatment^4^.

Growth failure, *e.g.* stunting (poor linear growth), is one of the most common manifestations of perinatally-acquired HIV infection, affecting up to 50% of children^5^. However, the impact on bone development in children is not well understood. A recent systematic review of bone health identified a high prevalence of low bone mass (when bone mass is two or more standard deviations [SD] below that expected for age) in children and adolescents living with HIV in high income countries (HICs), but found no studies from low income countries (LICs)^6^. Importantly, a one SD reduction in bone mass is associated with a doubling in childhood fracture risk, in otherwise healthy children^7^. In addition, adolescence is a crucial period of bone mass accrual, with peak bone mass (PBM) achieved at the end of skeletal maturation. Low PBM is a critical determinant of subsequent adult osteoporotic fracture risk^8^; a 10% reduction in PBM doubles fracture risk in adulthood^9,10^.

Muscle strength and bone strength are closely related; muscles exert forces on bone resulting in bone adaptation in size and strength^11^. HIV infection and consequent ill-health may result in reduced physical activity, which in turn may impair muscle strength and skeletal impact loading, impairing bone development^12–14^. Furthermore, ART may itself adversely affect bone and muscle health^15,16^. Few studies have compared muscle strength and function between HIV infected and uninfected children^13^. If greater muscle strength predicts higher bone mass, then physical activity interventions could potentially optimise musculoskeletal health.

The aim of this study was to determine the prevalence of low bone density among children with HIV (CWH) in Zimbabwe, and factors associated with low bone density, after accounting for body size. We further aimed to establish whether lean muscle mass and function are lower among CWH than uninfected peers, and whether muscle measures might explain any differences seen in size-adjusted bone density.

## Methods

A cross-sectional study was conducted between July 2016 and July 2017 at the paediatric HIV clinic at Harare Central Hospital, Zimbabwe. This is a public sector clinic that provides HIV care for more than 1,500 children. Children were eligible for the study if they were aged between 6 and 16 years, had been taking ART for at least 6 months, were not acutely unwell (no acute symptoms) and were not taking treatment for tuberculosis. Up to five eligible participants were consecutively recruited per day, restricted to this number due to logistical constraints. Children were convenience sampled within logistical and budgetary constraints. A comparison group of HIV-uninfected children in the same age group was recruited from seven primary care clinics that provided opt-out HIV testing and counselling to all attendees regardless of the reason for attendance, and that served the same catchment population as that of Harare Central Hospital. Children who tested HIV negative, were not acutely unwell and were not receiving treatment for tuberculosis were enrolled.

### Data collection

A nurse-administered questionnaire was used to collect socio-demographic data, clinical history including age at HIV diagnosis and ART initiation, ART regimen, history of menarche and voice breaking. Where possible, clinical history was confirmed with documentation within patient hand-held medical records. A standardized examination was performed including WHO staging of HIV infection and measurement of height and weight using SECA^®^ height board and electronic SECA^®^ weighing scales (Seca United Kingdom, Birmingham, England) and Tanner pubertal stage using standardised protocols and calibrated equipment. Hand grip strength in kilograms was measured using a Jamar hydraulic hand-held dynamometer (Patterson Medical, UK). Participants were seated with the shoulder at 0° to 10°, the elbow at 90° of flexion and the forearm positioned neutrally^17^. Three measurements were taken from each hand in alternation by trained staff, and the highest measurement from the six taken was used in analyses. Age and CD4 count at diagnosis were collected from hand held medical records and based on guardian report if no record was available.

### Dual-energy X-ray absorptiometry measurements

Dual X-ray absorptiometry (DXA) scans were performed by two trained radiographers using standard procedures on a Hologic QDR Wi densitometer (Hologic Inc., Bedford, MA, USA) with Apex Version 4.5 software for scan analysis. Daily calibration was conducted using a manufacturer provided spine phantom. Lumbar spine (LS) and Total body (TB) scans were performed from which BMC (Bone Mineral Content), TB lean mass, and TB fat mass were measured, and areal BMD (Bone Mineral Density) calculated (coefficients of variation <0.5%). DXA calculates two-dimensional (areal) BMD which is highly dependent upon bone size^18–20^ and is therefore unsuitable for use in growing children; DXA underestimates BMD when bones are small^21^. Therefore, size adjustment techniques, as recommended by the International Society for Clinical Densitometry (ISCD)^22^, were used to measure bone mineral apparent density (BMAD) Z-score at the lumbar spine and regression-based total-body less-head (TBLH) Bone Mineral Content (BMC) for lean mass adjusted for height (TBLH-BMC^LBM^) Z-score, using, in the absence of African reference datasets, the recommended age and gender specific reference data derived from growth curves for white children in the UK (the ALPHABET dataset: Amalgamated Reference Data for Size-Adjusted Bone Densitometry Measurements in Children and Young Adults)^23^. Total body-less-head (TBLH) BMC, which is considered one of the most accurate and reproducible methods in children, was measured^22^. Then Z-scores for TBLH-BMC adjusted for Lean Body Mass, fat mass and height (BMC^LBM^) were generated from ALPHABET regression-derived reference growth curves^23^. The head is excluded as it represents a greater proportion of skeletal mass in younger children and may obscure important skeletal deficits. Furthermore, the head is not weight-bearing and does not grow in response to factors like physical activity^22^. Low LS-BMAD and low TBLH-BMC^LBM^ were both defined as a Z-score ≤ −2, according to recommended clinical definitions^22^.

### Laboratory Investigations

Among CWH, HIV viral load was measured using COBAS Ampliprep/Taqman 48 Version 2.0 (Roche, Rotkreuz, Switzerland) and CD4 cell count was measured using an Alere PIMA CD4 (Waltham, Massachusetts, USA) machine.

### Ethical approval

Ethical approval was granted by the Medical Research Council of Zimbabwe (MRCZ/A/1856), the Harare Hospital Ethics Committee, the Biomedical Research and Training Institute Institutional Review Board (AP125) and the London School of Hygiene and Tropical Medicine Ethics Committee (8263). All guardians gave written consent, and participants gave written assent to participate in the study.

### Data analysis

Data were extracted from paper forms using optical character recognition software (Cardiff TELEFORM Intelligent Character, Version 10.7; Hewlett Packard, Palo Alto, California, USA). Data analysis was carried out using Stata v13 (StataCorp, College Station, Texas, USA).

Height-for-age and body mass-for-age Z-scores were calculated using the WHO reference standards^24^.

Continuous variables were summarised as mean (standard deviation: SD) and median (interquartile range: IQR), and categorical variables as counts (percentages). Multivariable linear regression models were used to examine the differences in bone and muscle measures between participants with and without HIV infection. Continuous outcome variables were standardized for regression analyses, so that differences are presented, with 95% confidence intervals (95% CI), as the number of SDs difference between HIV infected and uninfected participants. Serial adjustments to differences in bone and muscle measures were decided *a priori;* model 1: adjusted for age and sex, model 2: adjusted for age, sex, height (categorised), model 4: adjusted for age, sex, height (categorised by 10cm increments to account for any non-linearity), and puberty. As Hologic derived TBLH-BMC^LBM^ takes account of lean mass and height, these were not adjusted for further in TBLH-BMC^LBM^ models. Modification of the effect of age at ART initiation by tenofovir use on bone outcomes, of lean mass and muscle function by HIV status on bone outcomes, and of age at the time of DXA by HIV status on bone and muscle outcomes, were investigated by fitting appropriate interaction terms to linear regression models. In addition, we calculated correlation coefficients for figures comparing two linear variables.

## Results

A total of 97 CWH and 77 uninfected participants were recruited, of whom 52% were female. Participants with HIV were older than those uninfected, mean (SD) 12.7 (2.5) *vs*. 10.0 (2.9) years, and hence were taller and heavier (Table 1). A high proportion of CWH were stunted compared to those uninfected (32% *vs*. 5%, *p*<0.001). All CWH were infected perinatally, and 22% of HIV-uninfected individuals reported having a mother with HIV.

**Table 1:** Characteristics of study participants by HIV status

|  | HIV-positive<br>(n=97) | HIV-negative<br>(n=77) | *p* value |
| --- | --- | --- | --- |
| **General characteristics** |  |  |  |
| *Mean (SD)* |  |  |  |
| Age at DXA scan, years | 12.7 (2.5) | 10.0 (2.9) | <0.001 |
| Height, cm | 141.5 (11.4) | 134.0 (16.6) | <0.001 |
| Weight, kg | 35.2 (9.5) | 31.1 (10.5) | 0.007 |
| BMI, kg/m<sup>2</sup> | 17.3 (2.6) | 16.8 (1.9) | 0.175 |
| *N (%)* |  |  |  |
| Gender, female | 51 (52.6) | 40 (52.0) | 0.934 |
| Puberty (menarche or voice broken)<sup>a</sup> | 25 (26.0)<sup>b</sup> | 11 (14.3) | 0.058 |
| Mother known to be HIV infected | 97 (100) | 17 (22.1) | <0.001 |
| Stunted (HFA Z-score ≤-2) | 31 (32.0) | 4 (5.2) | <0.001 |
| Underweight (BFA Z-score ≤-2) | 14 (14.4) | 0 (0.0) | 0.001 |
| **DXA and grip strength characteristics** |  |  |  |
| *Mean (SD)* |  |  |  |
| Maximal handgrip strength, kg | 21.0 (8.2)<sup>c</sup> | 16.2 (6.5)<sup>d</sup> | <0.001 |
| TB Lean mass, kg | 26.2 (6.3)<sup>e</sup> | 23.0 (7.8) | 0.003 |
| TB Fat mass, kg | 9.1 (4.6)<sup>e</sup> | 8.6 (3.7) | 0.397 |
| TBLH-BMC, g | 896.0 (262.7)<sup>e</sup> | 789.0 (333.4) | 0.019 |
| TBLH-BMC<sup>LBM</sup> Z-score | -0.643 (1.189) | -0.314 (0.882) | 0.045 |
| L spine BMC, g | 28.4 (8.9) | 24.9 (11.4) | 0.023 |
| L spine BMAD, g/cm<sup>2</sup> | 0.196 (0.037) | 0.195 (0.032) | 0.801 |
| L spine BMAD Z-score | -0.742 (1.295)<sup>e</sup> | -0.093 (0.886) | <0.001 |
| *N (%)* |  |  |  |
| Low TBLH-BMC<sup>LBM</sup> (Z-score ≤-2) | 12 (12.5)<sup>e</sup> | 2 (2.6) | 0.018 |
| Low L Spine BMAD (Z-score ≤-2) | 14 (14.6)<sup>e</sup> | 1 (1.3) | 0.002 |

BMI, Body Mass Index; HFA, Height for Age; BFA, Body Mass for Age; SD, Standard Deviation; DXA, Dual energy X-ray Absorptiometry; TB, Total Body; TBLH, Total-Body Less Head; BMC, Bone Mineral Content; TBLH-BMC<sup>LBM</sup>; total-body less-head Bone Mineral Content for lean mass adjusted for height; BMAD, Bone Mineral Apparent Density; L spine, Lumbar Spine.
<sup>a</sup> mean age at puberty 11.9 (2.6) years for CWH and 9.5 (2.9) years for children without HIV. 16 girls with HIV had reached menarche at mean [SD] age 13.5 [2.5] years, compared with 7 girls without HIV 13.2 [1.4] years.
<sup>b</sup>n=96 CWH with data, <sup>c</sup>n=85. <sup>d</sup>n=71. <sup>e</sup>n=96.
Students t test for continuous data and $\chi^2$ test for categorical data (*i.e.* unadjusted)

The 97 CWH had a median age at diagnosis of 4 years (IQR 2-7) (Table 2). The median age at initiation of ART was 5.9 years (IQR 3.2-8.4) and 44 (46%) reported past or current tenofovir use. At enrolment, 79% were virologically supressed (HIV viral load <400 copies/ml) and the median CD4 count was 733 (IQR 473-983) cells/mm^3^. Only one child with HIV reported a prior history of an arm fracture.

**Table 2:**
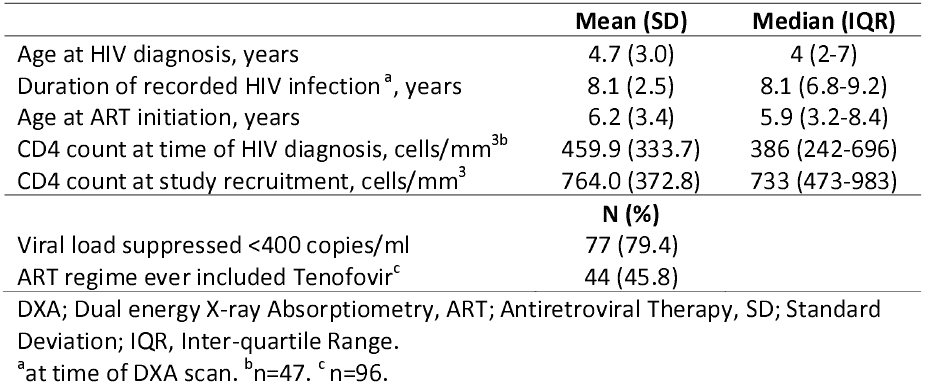
HIV-specific summary characteristics amongst the 97 children with HIV

### Bone density

Low LS-BMAD (*i.e*. a Z-score ≤-2) was more common among CWH than those uninfected, 15% *vs*. 1% (*p*=0.002); the same was true for low TBLH-BMC^LBM^ found in 13% *vs*. 3%, *p*=0.018 (Table 1). Among CWH, there was no association between stunting, defined as a height-for-age (HFA) Z-score <-2, and of low BMAD at the lumbar spine (*p* value= 0.36) or of low TBLH-BMC^LBM^(*p* value= 0.81).

Initially, in unadjusted analyses, CWH appeared to have higher lumbar spine BMC than uninfected individuals (Table 1), but when comparing BMAD, which takes account of body size, this difference was no longer observed (Table 1). However, when then comparing BMAD Z-scores, which standardises for age and gender, the LS-BMAD Z-scores were significantly lower in CWH compared to uninfected children (mean −0.742 *vs*. −0.093, *p*<0.001) (Table1). Similar patterns were seen in the TBLH after taking account of body size and after standardisation for age and gender, with a mean TBLH-BMC^LBM^ Z-score of −0.643 in CWH compared to Z-score −0.314 in those uninfected, *p*=0.045. After adjustment for age, sex, height and puberty, a difference was still seen in LS-BMAD between CWH and uninfected children, such that the mean difference was 0.33 SDs (95% CI −0.01, 0.67; *p*=0.055) lower in CWH compared to those without HIV, whilst no clear difference in TBLH-BMC^LBM^ [0.26 SDs (95% CI −0.09, 0.61; *p*=0.148)] was observed between CWH and those without HIV.

In CWH, there was a concerning inverse association between age and LS-BMAD Z-Score, and to a lesser extent TBLH-BMC^LBM^ Z-Score (Figure 1, unadjusted). This contrasted with the stable LS-BMAD and TBLH-BMC^LBM^ Z-Scores seen across all ages among children without HIV. This suggests that older CWH are more likely to have a low LS-BMAD in comparison with their peers without HIV. In support of this, there was modification of the effect of age on LS-BMAD Z-score by HIV status (interaction *p*=0.03).

**Figure 1:**
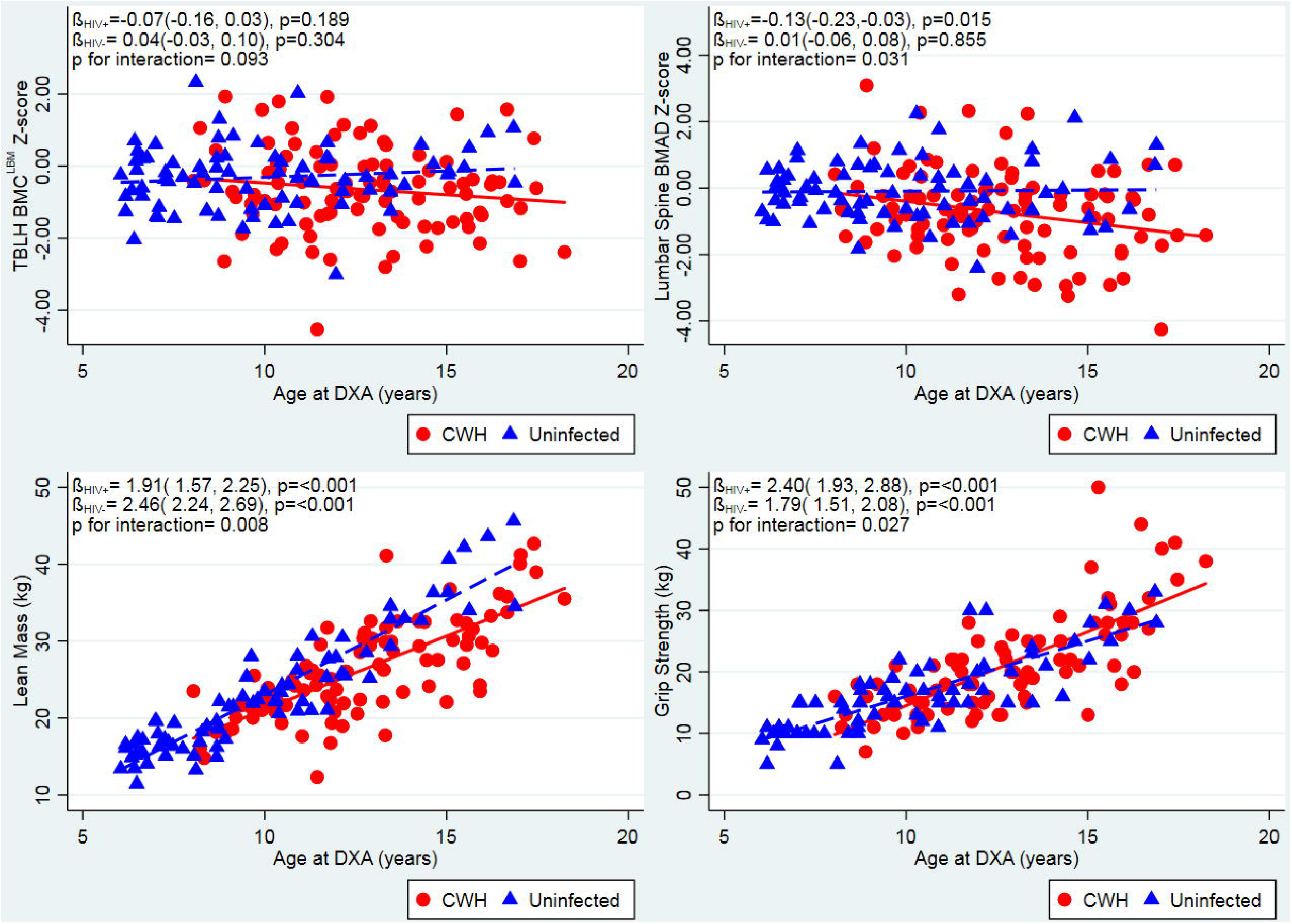
Unadjusted relationships between age and TBLH-BMC^LBM^ and LS BMAD Z-scores, lean muscle mass, grip strength, stratified by HIV status. Red unbroken line and squares indicate children with HIV (CWH). Blue dotted line and triangles indicate children without HIV infection. DXA, Dual energy X-ray Absorptiometry; TBLH-BMC^LBM^; total-body less-head Bone Mineral Content for lean mass adjusted for height; BMAD, Bone Mineral Apparent Density.

### Bone density and HIV treatment

There was a strong association between age at ART initiation and both TBLH-BMC^LBM^ Z-score and LS-BMAD Z-score, such that for each year ART initiation was delayed, there was a 0.08 SD reduction in TBLH-BMC^LBM^ and 0.13 reduction in LS-BMAD (Figure 2), although correlations were weaker. This means that those children starting ART after the age of 8 years had on average, at least a 1 SD reduction in LS-BMAD compared with the reference population. There was no modification of this effect by the use of tenofovir as part of the ART regimen (*p*=0.37), but numbers were small (Supplementary Figure 1). Those who took tenofovir had a significantly lower LS-BMAD Z-scores than those who had not received tenofovir (mean [SD] −1.14 [1.28] *vs*. −0.44 [1.23], *p*=0.008); a less marked difference was seen for TBLH-BMC^LBM^ Z-Score (−0.90 [1.02], *vs*. −0.46 [1.28], *p*=0.072).

**Figure 2:**
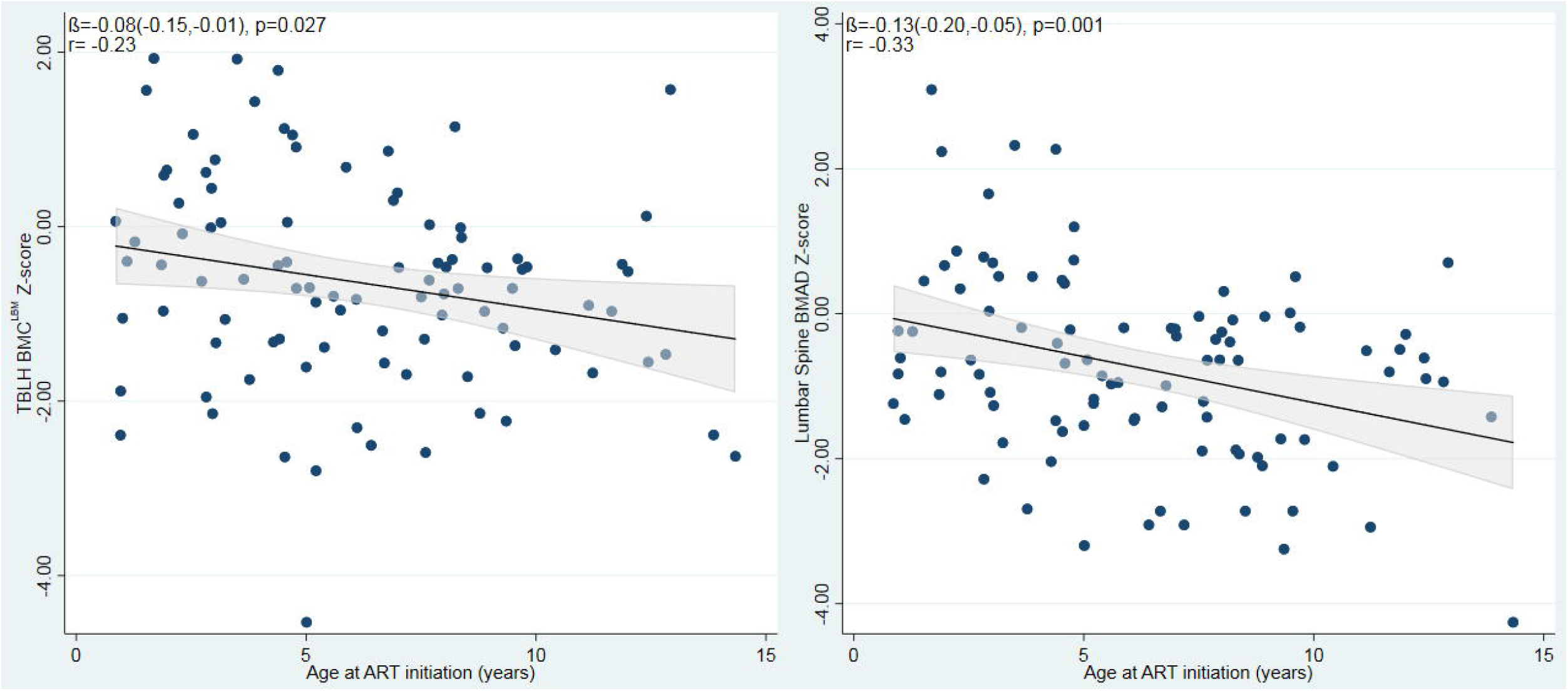
Unadjusted relationships between age at ART initiation and TBLH-BMC^LBM^ and Lumbar spine BMAD Z-scores among children with HIV. ART; Antiretroviral Therapy; TBLH-BMC^LBM^; total-body less-head Bone Mineral Content for lean mass adjusted for height; BMAD, Bone Mineral Apparent Density.

### Muscle Mass & Function

Age was positively associated with lean mass and with muscle function (grip strength), in both CWH and children without HIV, although uninfected children demonstrated a stronger association between age and lean mass, than did CWH (interaction *p*=0.008) (Figure 1). The greater body weight measured among CWH compared to those children without HIV was explained by a higher relative proportion of lean mass, rather than increased fat mass (Table 1). Correspondingly, muscle function, measured as maximal grip strength, was found to be greater among CWH than those uninfected; however, after adjusting for age and sex, no difference in muscle function was observed (Table 3). After similar adjustment, CWH had a lower lean mass than those who were uninfected, but this was explained by differences in height and pubertal stage between the two groups (Table 3). No clear relationships were seen between either lean mass or muscle function and TBLH-BMC^LBM^ or LS-BMAD Z-scores, in participants with or without HIV (Supplementary Figure 2).

**Table 3:** Differences in Lumbar Spine BMAD, TBLH-BMC^LBM^, muscle mass and muscle function between children with and without HIV infection, after serial adjustments

|  | N | Mean difference<br>(95% CI) | p value |
| --- | --- | --- | --- |
| *TBLH-BMC<sup>LBM</sup>* |  |  |  |
| Model 1: age, sex | 172 | -0.30 (-0.64, 0.04) | 0.088 |
| Model 3*: age, sex, puberty | 172 | -0.26 (-0.61, 0.09) | 0.148 |
| *Lumbar spine BMAD* |  |  |  |
| Model 1: age, sex | 172 | -0.41 (-0.73, -0.09) | 0.011 |
| Model 2: age, sex, height | 172 | -0.39 (-0.71, -0.06) | 0.022 |
| Model 3: age, sex, height, puberty | 172 | -0.33 (-0.67, 0.01) | 0.055 |
| *Lean Muscle Mass* |  |  |  |
| Model 1: age, sex | 172 | -0.38 (-0.56, -0.20) | <0.001 |
| Model 2: age, sex, height | 172 | -0.16 (-0.29, -0.03) | 0.019 |
| Model 3: age, sex, height, puberty | 172 | -0.11 (-0.24, 0.03) | 0.111 |
| *Muscle grip strength* |  |  |  |
| Model 1: age, sex | 154 | -0.10 (-0.31, 0.11) | 0.353 |
| Model 2: age, sex, height | 154 | -0.01 (-0.22, 0.20) | 0.924 |
| Model 3: age, sex, height, puberty | 154 | 0.05 (-0.16, 0.27) | 0.620 |

BMAD; Bone Mineral Apparent density, TBLH-BMC<sup>LBM</sup>; total-body less-head Bone Mineral Content for lean mass adjusted for height. CI; Confidence interval.
Model 1: adjusted for age and gender. Model 2: model 1 & height (as category). Model 3: model 2 & puberty. Model 3\*: adjusted for age, gender and puberty. Mean differences are shown in SD units, negative Betas (mean difference values) indicate a lower measured outcome in the group with HIV infection.

## Discussion

Our study showed that low size-adjusted bone density is common among CWH, the majority of whom were pre-pubertal. This low bone density was observed in both trabecular-rich skeletal sites (*i.e*. 15% having low lumbar spine BMAD) and cortical-rich sites *(i.e.* 13% having low TBLH-BMC^LBM^). Importantly, the age at ART initiation was a key predictor of lumbar spine bone density, such that children starting ART after the age of 8 years had on average, at least a 1 SD reduction in lumbar spine BMAD compared with the reference population. This is a clinically important effect size, as it is thought that a 1 SD reduction in bone density doubles both childhood and, if sustained, future adult fracture risk^7,25^. Given the recent report that current median age of ART initiation in Sub-Saharan Africa is 7.9 years^26^, the extrapolation that potentially half of such children may be at significant increased fracture risk is of concern. There was also evidence of lower bone density, particularly in trabecular-rich skeletal sites, in those children treated with tenofovir as part of their ART regimen. In Zimbabwe tenofovir forms part of the first-line ART regimen in children aged 12 years or older and/or who weigh 35kgs or more. Hence, our findings from a relatively young population, with a relatively short duration of tenofovir exposure, may underestimate any effect of tenofovir on the growing skeleton. These findings are consistent with several studies in adults^27^; however, data from children have been less consistent^28^ and longitudinal data in larger populations, including those in Sub-Saharan Africa, are needed to determine whether this association is transient or persists after treatment initiation.

Our findings show that stunting is not a proxy for low size-adjusted bone density. Adolescent skeletal growth is not linearly and structurally uniform; peak height velocity precedes peak bone content accrual^29^, yet how bone is accrued in the context of HIV infection with associated delayed puberty, and hence skeletal maturation^30^, is poorly understood. Despite CWH being on average 2.7 years older than those uninfected, only 14% more CWH had entered puberty. HIV infection is an established cause of pubertal delay and it is possible that some of the effect of HIV infection on size-adjusted bone density may be mediated through this mechanism^31,32^.

After accounting for body size and pubertal stage, no differences were seen in muscle mass or function between those with and without HIV infection. Furthermore, neither muscle mass or function were associated with bone density, suggesting that muscle is not affected in the same way as bone in the context of longstanding paediatric HIV infection. A small study of 15 Puerto Rican CWH were also found to have similar lower limb muscle strength to uninfected controls^33^. Although, a more recent Canadian study which assessed lower limb muscle function by jumping mechanography, in a population 2 years older than ours (n=35), did find an albeit small difference in lower limb muscle power^13^. Whether power rather than strength, or lower limb rather than upper limb function, may be affected by HIV infection remains to be determined.

The Children with HIV Early Antiretroviral Therapy (CHER) trial showed substantially reduced mortality in infants with immediate ART initiation leading to recommendations that all infants with HIV should be treated with ART following diagnosis^34^. There has been no such benefit demonstrated in older children^35,36^ and until recently, guidelines for treatment eligibility for older children and adolescents were the same as those for adults, based on disease or immunological stage^37,38^. The recent START and TEMPRANO trials both demonstrated that early initiation of ART reduced the risk of severe illnesses^39,40^; these trials were conducted in adults only and excluded older children and adolescents. Our findings add strong support to the recent WHO treatment guidelines which recommend prompt ART initiation following diagnosis of HIV infection regardless of clinical or immunological stage of infection in children^38^. While these guidelines are a positive step towards improvement of health of CWH, there likely remains a cohort of children in Sub-Saharan Africa who are missed by programmes to prevent mother-to-child transmission including early infant diagnosis and who will therefore be diagnosed in later childhood^41^. Furthermore, children progressing through adolescence are at high risk of poor adherence and therefore inadequate viral suppression, which may put their bone health at risk^42^. It remains to be seen whether improved ART roll-out will ameliorate this risk, or whether there will be a higher than expected fracture burden in the region in the future.

Our study highlights the importance of using the ‘gold-standard’ size-adjustment methodology when conducting studies of bone density in the growing skeleton, particularly in the context of HIV where stunting is so common. Other strengths of this study are the prospective and (based on symptoms) unselected recruitment of participants. Until now, DXA-based studies in CWH in Sub-Saharan Africa have been limited to South Africa^43^. DXA capacity is severely limited across the region^44^ so that such studies are logistically challenging. Our study has limitations, including the cross-sectional nature of the study design which prevents causal inference, and the older age of the infected compared to uninfected participants which explain the height and weight differences seen, although our analyses enabled suitable adjustment. We lacked hand/wrist radiographs from which to derive bone age as a more accurate measure of skeletal maturation than chronological age adjusted for Tanner stage. We lacked data to permit analysis of differing ART regimes and durations and their associations with weight, height and bone measurements. Notably, 22% of our uninfected children reported maternal HIV infection, hence these children may have been exposed but uninfected. It is unclear whether such children are at risk of impaired skeletal development, if so, this may have led us to underestimate the true magnitude of difference between CWH and uninfected (and unexposed) children. Furthermore, there is a global lack of bone density reference data for child/adolescent populations in Africa, which meant that following International Society for Clinical Densitometry (ISCD) guidance^22^, we used the recommended age and gender specific reference data derived from white children in the UK^23^. Future studies are needed to establish normative data for African populations.

In conclusion, perinatally-acquired HIV infection is associated with both stunting and low lumbar spine bone density (adjusted for size) in Zimbabwean children, but importantly these conditions appear to be independent of one another. Such low bone density has been associated with substantially increased fracture risk in other populations. Our findings suggest delays in initiating ART contribute to lower bone density in CWH, and our findings support the current recommendations to initiate ART regardless of disease or immunological stage. Longitudinal data are needed from high HIV prevalence settings in Sub-Saharan Africa to establish how size-adjusted bone density changes through pubertal growth and following treatment of HIV infection, together with future fracture incidence studies.

## Acknowledgements

CLG is funded by Arthritis Research UK (grant ref 20000), AH is funded by Wellcome Trust (grant ref 20378/Z/16/Z), RR is funded by Wellcome Trust (grant ref 206764/Z/17/Z), RAF is funded by Wellcome Trust (grant no 095878/Z/11Z and the Norwegian Research Council through GLOBVAC. Reference data were collected as part of a Linda Edwards Memorial Studentship (KW PI) and funded through Central Manchester Foundation NHS Trust endowments. We would like to thank Abel Karera for his assistance acquiring the DXA scans. We further thank all our study participants.

## Author’s contributions

RAF designed the study. CLG and RR drafted the manuscript. AH performed the statistical analysis. EM, GM, TB study oversight, data collection and preliminary analysis. LS reviewed study protocol. CMK performed the DXA scans. NC derived size-adjusted DXA data. KW was PI of the study which provided control data and advised on data analysis. All authors provided feedback on the draft manuscript and approved the final manuscript.

---

**Supplementary Figure 1:** Unadjusted relationships between age at ART initiation and TBLH-BMC^LBM^ and Lumbar spine BMAD Z-scores, stratified by the use of Tenofovir, among children with HIV. Red unbroken line and squares indicate children with HIV and a history of Tenofovir use. Blue dotted line and triangles indicate children with HIV without a history of Tenofovir use. ART; Antiretroviral Therapy; TBLH-BMC^LBM^; total-body less-head Bone Mineral Content for lean mass adjusted for height; BMAD, Bone Mineral Apparent Density.

**Supplementary Figure 2:** Unadjusted relationships between muscle mass and function and TBLH-BMC^LBM^ and Lumbar spine BMAD Z-scores, stratified by HIV status. Red unbroken line and squares indicate children with HIV. Blue dotted line and triangles indicate children without HIV. TBLH-BMC^LBM^; total-body less-head Bone Mineral Content for lean mass adjusted for height; BMAD, Bone Mineral Apparent Density.

