## Supplementary figures and images for "Older age at Initiation of Antiretroviral Therapy Predicts Low Bone Mineral Density in Children with perinatally-infected HIV in Zimbabwe"

### Supplementary Figure 1

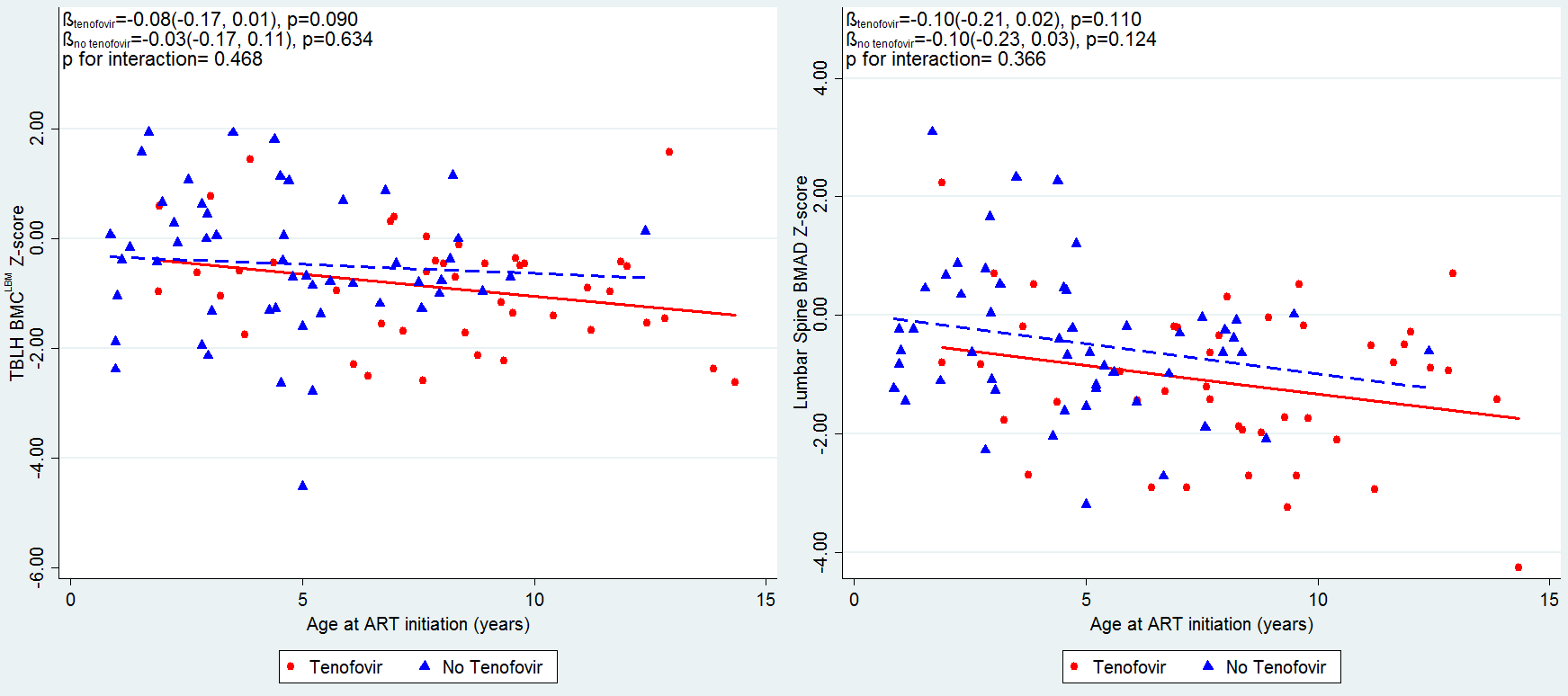

### Supplementary Figure 2

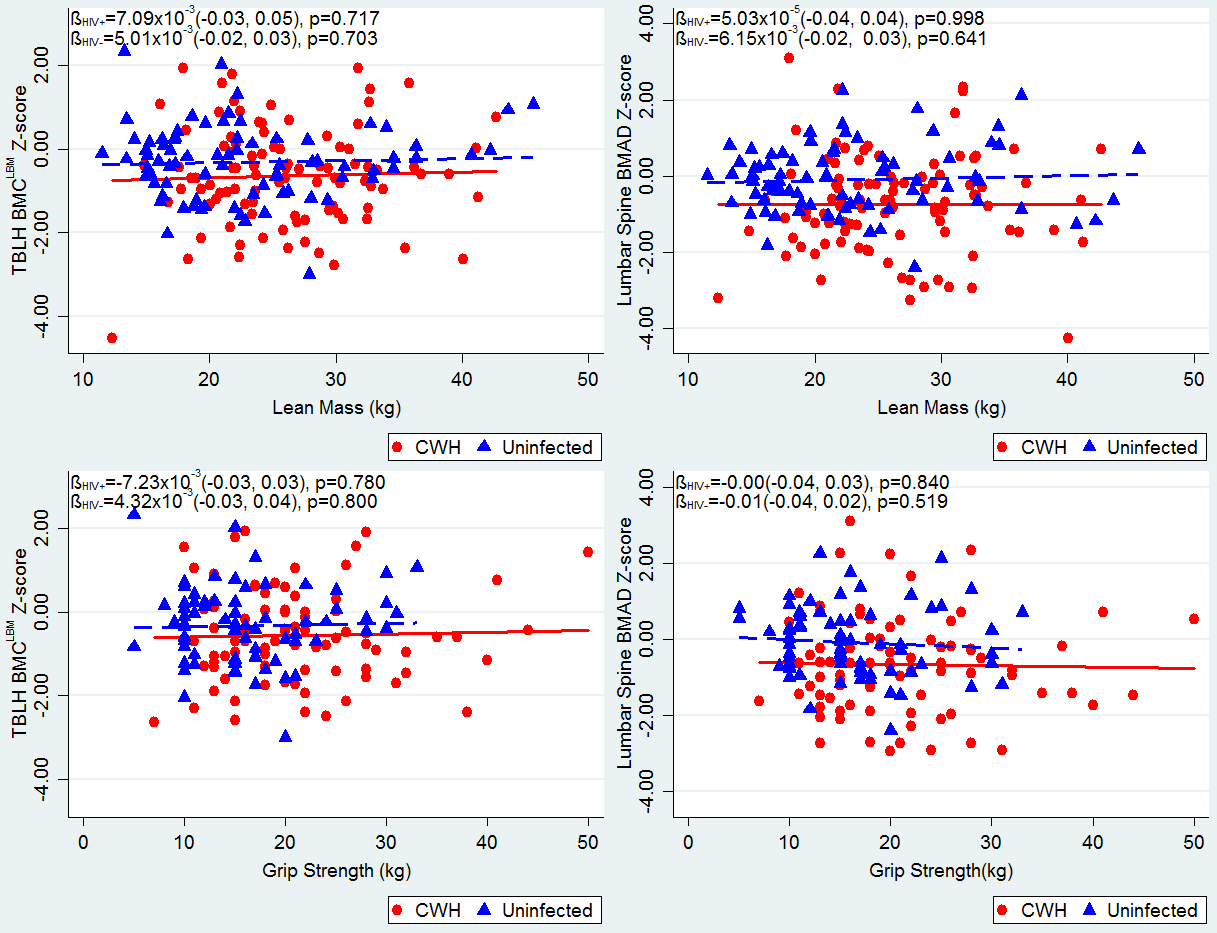
